# Environmental and Historical Determinants of African Horse Sickness: Insights from Predictive Modeling

**DOI:** 10.1101/2024.02.20.581150

**Authors:** KwangHyok Kim, TianGang Xu, Arivizhivendhan Kannan Villalan, TianYing Chi, XiaoJing Yu, MyongIl Jin, RenNa Wu, GuanYing Ni, ShiFeng Sui, ZhiLiang Wang, XiaoLong Wang

## Abstract

This study marks a pioneering effort in analyzing the global epidemiological patterns of African Horse Sickness (AHS) across different regions. By employing predictive modeling with a comprehensive set of environmental variables, we uncovered overarching global patterns in AHS dynamics, a first in this field. Our analysis revealed significant regional differences influenced by specific climatic conditions, highlighting the disease’s complexity. The study also identifies new high-risk areas for AHS, underscoring the necessity for regionally tailored disease management strategies. Despite some limitations, such as the exclusion of wild equine data, this research offers critical insights for global AHS intervention and prevention, setting a path for future research incorporating broader datasets and socio-economic factors.

**Author Summary:** AHS presents a significant challenge to the global equine industry, impacting both health and economic aspects. Our study highlights the profound effect of climate change, particularly the frequency of extreme climatic events including temperature and humidity variations, on the transmission dynamics of diseases like AHS. In our research, we focused on overcoming the challenges associated with identifying key environmental factors and determining the appropriate geographic scale for a comprehensive global understanding of AHS. Our aim was to bridge existing knowledge gaps and elucidate the fundamental principles governing AHS transmission. This study establishes a solid foundation for understanding the intricate dynamics of AHS and offers practical pathways for conservation efforts. It emphasizes the urgent need for environmentally conscious strategies to protect horse populations and the industries dependent on them. By highlighting the relationship between environmental factors, vector presence, and AHS transmission, our research underscores the importance of a holistic approach to disease mitigation. In conclusion, the findings of our study not only contribute to the scientific understanding of AHS but also serve as a guide for policymakers and practitioners in developing effective strategies for disease management and prevention, tailored to the specific environmental conditions and challenges faced in different regions around the world.

## 1. Introduction

African Horse Sickness (AHS), a viral disease affecting equine species, is primarily transmitted by arthropods and significantly influenced by environmental conditions (1–3) as well as human activities (4). The complexity of its transmission dynamics, shaped by both natural and anthropogenic factors, sets the stage for a comprehensive exploration of its historical spread and ecological implications. In the context of ongoing global warming and changing land-use patterns, it is anticipated that the distribution and prevalence of AHS will undergo notable alterations (1, 2). The African Horse Sickness Virus (AHSV), classified within the *Orbivirus* genus, encompasses nine distinct serotypes. Among equine species, horses (*Equus ferus*) and donkeys from Europe and Asia *(E. hemionus*) exhibit the highest susceptibility, with mortality rates reaching up to 90%. In contrast, African donkeys (*E. africanus*) and various zebra species (*E. quagga, E. grevyi*, and *E. zebra*) often remain asymptomatic, thereby serving as natural reservoirs for AHSV (5).

Tracing the historical trajectory of AHS reveals its expanding geographic scope. First documented in 1569 in Central and Eastern Africa (6), the disease has since marked its presence\ across diverse regions including the Middle East, Western Asia, North Africa, the Mediterranean basin, and more recently, Southeast Asia with incidents in Thailand and Malaysia in 2020 (7–14). This historical perspective lays a foundation for understanding the disease’s current distribution and potential vectors. Contrary to some zoonotic diseases, AHSV transmission does not occur through direct contact between animals. Instead, it primarily depends on hematopoietic arthropod vectors. The *Culicoides* species, including *C. imicola, C. bolitinos, C. variipennis*, and *C. brevitarsis*, are the principal vectors facilitating regular transmission. Other vectors like mosquitoes (*Aedes aegypti, Anopheles stephensi*, and *Culex pipiens*), ticks (*Hyalomma dromedarii* and *Rhipicephalus sanguineus*), and biting flies (*Stomoxys calcitrans* and *Tabanus* spp.) contribute to intermittent transmission modes (15). The geographical expansion of AHS correlates closely with the widening of the AHSV suitability zone, a phenomenon largely attributable to environmental factors influencing the distribution and behavior of AHSV-associated species.

Building upon the historical context, it is crucial to examine how environmental variables such as temperature and humidity influence the spread of AHS. These factors not only affect the virus’s survival but also the behavior and population dynamics of the host and vectors, thereby playing a pivotal role in the epidemiology of the disease. Factors such as temperature, humidity, topography, and land cover have been extensively studied, revealing their significant impact on the virus’s viability and the behavioral patterns of both the host species and the vectors. Research indicates that these environmental conditions directly influence the survival duration of the virus and affect the range of activities of the host and vector species (16, 17). This suggests a complex interplay between the ecological landscape and the epidemiology of AHS, underscoring the need for a comprehensive understanding of environmental influences on the disease’s transmission dynamics.

Further delving into specific environmental factors, temperature and relative humidity emerge as key determinants in the transmission of AHSV. Their impact on the lifecycle and behavior of vectors such as *Culicoides* and mosquitoes highlights the intricate relationship between climatic conditions and disease dynamics. Specifically, temperature governs the rate of development in immature vectors and markedly affects their behavior and lifespan. Within certain temperature ranges, an increase in temperature results in a higher egg production rate, faster larval development, and an elongated period of biting activity in female *Culicoides* (18). Conversely, extremely low temperatures inhibit *Culicoides* development and activity, whereas excessively high temperatures lead to increased mortality and reduced lifespans (19). Similar temperature-dependent behavioral patterns are observed in mosquitoes (*Aedes aegypti, Anopheles stephensi*, and *Culex pipiens*) (20–23), camel ticks (*Hyalomma dromedarii*), brown dog ticks (*Rhipicephalus sanguineus*) (24, 25), and biting flies from the *Stomoxys* and *Tabanus* genera (26, 27).

In addition to temperature, precipitation plays a critical role in shaping the transmission patterns of AHS. This aspect of environmental influence provides insights into the breeding habits of vectors and their consequent impact on the spread of the disease, illuminating the multifaceted nature of AHS transmission. Precipitation exerts a substantial influence on the development stages of immature vectors and the activity and dispersal patterns of adult vectors, primarily through its effects on atmospheric and soil humidity levels. In certain regions of South Africa, notably higher populations of *C. imicola* were recorded during years with abundant rainfall (28). Furthermore, significant outbreaks of AHS in South Africa have been closely associated with periods of heavy rainfall followed by drought conditions (29). These observations suggest a strong correlation between the presence of favorable breeding sites, supporting the immature development of *C. imicola*, and the incidence of AHS outbreaks. The breeding environments for mosquito larvae, such as water-filled containers, as well as the flight patterns and egg-to-larva transition process in biting flies, are also directly influenced by specific rainfall patterns (30). Collectively, these factors may significantly contribute to the spread of AHS through intermittent transmission modes, resulting in a cumulative effect. However, it is important to note that excessive rainfall can negatively impact AHS transmission. For instance, heavy rains can lead to floods that wash away larvae, resulting in a decline in vector populations (31).

Beyond climatic factors, land cover and altitude are also instrumental in understanding the spread of AHS (17, 32, 33). These elements contribute to the creation of suitable habitats for vectors, thereby influencing the distribution and intensity of AHS outbreaks. Research has pinpointed various land cover types, such as urban areas, croplands, and herbaceous tree shrubs, as conducive environments for AHS vectors, with these habitats being significantly influenced by alterations in altitude and land cover (34–36). Additionally, the transportation and trade networks of horses are considered crucial in the dissemination of AHS (37, 38), highlighting the importance of stringent quarantine measures in horse movement. However, monitoring equine populations presents considerable challenges, and obtaining a thorough and accurate assessment of equine movement patterns is a daunting endeavor. Furthermore, given the observed alignment of AHS spread with windborne movements of infected midges, researchers have proposed the potential for AHS transmission via wind dispersion. Recent studies have incorporated wind speed as a variable in modeling the spatiotemporal distribution of AHS and *C. imicola* (33, 39, 40). Despite these advancements, the optimization of *Culicoides* spp. dispersal through specific wind characteristics and its potential variations across regions and seasons remains an area of ongoing research and uncertainty.

In light of these diverse environmental influences (17, 33, 41), our study aims to integrate these varied environmental factors into a cohesive model. By employing a global-scale approach and focusing on variable identification, we endeavor to elucidate the general laws governing the spread of AHS and address the geographical variability in prediction accuracy. Our objective is to discern the overarching patterns in AHS distribution by identifying key variables and addressing the challenges posed by geographic scale selection in predictive modeling. To this end, we have utilized the MaxEnt model to define the ecological niches of AHS and its primary and intermittent transmission vectors. To the best of our knowledge, this study represents a pioneering effort to model the global dynamic laws governing AHSV distribution, thereby contributing novel insights to the field.

## 2. Materials and Methods

### 2.1. Data Collection

#### 2.1.1. AHS Occurrence Data

Location data pertaining to AHS were obtained from two primary sources: the Global Animal Disease Information System (EMPRES-i) website and reports published by the WOAH. This data forms the basis for identifying the geographical distribution of AHS.

#### 2.1.2. AHSV Vector Data

The presence of AHSV vector species was compiled from a variety of sources. This includes a comprehensive review of existing literature accessible through academic databases such as Web of Science, Science Direct, and PubMed. Additionally, data was sourced from the Global Biodiversity Information Facility (GBIF) database (available at: https://www.gbif.org/). To ensure the accuracy and relevance of this data, any records that lacked specific geographic details were excluded from our analysis.

#### 2.1.3. Environmental Predictors and Horse Data

Our modeling approach integrated various environmental factors that potentially influence AHS distribution. These factors included climate conditions, terrain features, land cover types, and the distribution of horse populations. The sources for this data were globally recognized databases, ensuring a high standard of data quality and reliability. Prior to integration into our models, all spatial data were pre-processed using ArcGIS software, version 10.6. This pre-processing involved standardizing and resampling the data to achieve a uniform resolution of 30 arc seconds, facilitating more accurate and consistent analyses.

### 2.2. Climate-Based Zonal Segregation

Various climate patterns have a significant impact on the distribution of species and biodiversity gradients within a given area. Climate factors are widely known to be the fundamental drivers of AHS occurrence and there are noticeable disparities in key climate variables associated with AHS across regions with different climate characteristics (42). As a solution to address the inconsistencies in the major environmental factors affecting AHS and their level of influence attributable to the chosen geographical scale, using the Köppen climate map, regions were categorized into tropical, arid, temperate, continental, and polar/alpine climates. For modeling efficiency, each climate zone was further segmented by continent.

### 2.3. AHS ecological niche modeling

Spatial autocorrelation was addressed by ensuring a minimum 10 km distance between recorded AHS locations using SDM Toolbox v1.1c (17). In the identification of crucial variables associated with AHS, careful consideration was given to the process of reducing multicollinearity. This was done to prevent any distortion in the model caused by correlations between variables. Principal component analysis (PCA) and variance inflation factor (VIF) analysis were employed to resolve the problem of multicollinearity among predictors (43). Frist, major climate predictors were selected in SPSS 2.2 using eigenvalues larger than 1.0 and the Scree Plot criterion or “broken stick” stopping rule. Next, MaxEnt model analysis was performed to remove climate variables with high standard deviation (SD) and low contribution rate based on visual observation of the response curve and percent contribution. Finally, VIF analysis was performed on selected climate variables and other variables to assess collinearity between environmental variables. If the VIF value was <10, it was assessed as low multicollinearity and input was allowed. The regions where AHS occurrence was verified were separated according to their climate zones, modelled individually, and subsequently amalgamated. The result of this process is a comprehensive AHS suitability map. Model reliability was gauged using the Area Under the Curve (AUC), considering AUC>0.8 as an indicator of a well-fitted model.

### 2.4. AHSV Vectors Niche Modeling

In our research, we extended our focus beyond the commonly studied *Culicoides* spp. to include a broader range of potential carriers. This expansion encompasses mosquitoes, ticks, and biting flies, all recognized by the WOAH as vectors capable of intermittent transmission of AHS (15). By incorporating vectors linked to both primary and intermittent transmission modes of AHSV, our model achieves a more thorough and nuanced understanding of the potential risk areas for AHS.

The methodology employed to minimize spatial autocorrelation and to execute Principal Component Analysis (PCA) and Variance Inflation Factor (VIF) analyses is consistent with that used in AHS ecological niche modeling. Our approach involved confirming the presence of vector species based on climate zones, with each zone being modeled individually. We then integrated the modeling outcomes for each climate zone to create suitability maps for individual vector species. These maps were combined to form an overarching vector suitability map that includes all species under consideration. A model was considered well-fitted if its Area Under the Curve (AUC) value exceeded 0.8.

### 2.5. Identifying High-Risk Zones for AHS by Integrated Modeling Approach

To determine the likelihood of AHS occurrence in different regions, we employed an integrated modeling approach. Initially, vector suitability maps were overlaid with horse density data. This was followed by the addition of the AHS suitability map, culminating in the generation of a comprehensive AHS risk map. The risk levels were stratified into four categories: low (< 0.25), medium (0.25-0.5), high (0.5-0.75), and very high (> 0.75). Fig. 1 in the manuscript illustrates a flow chart detailing our modeling approach.

**Fig. 1.**
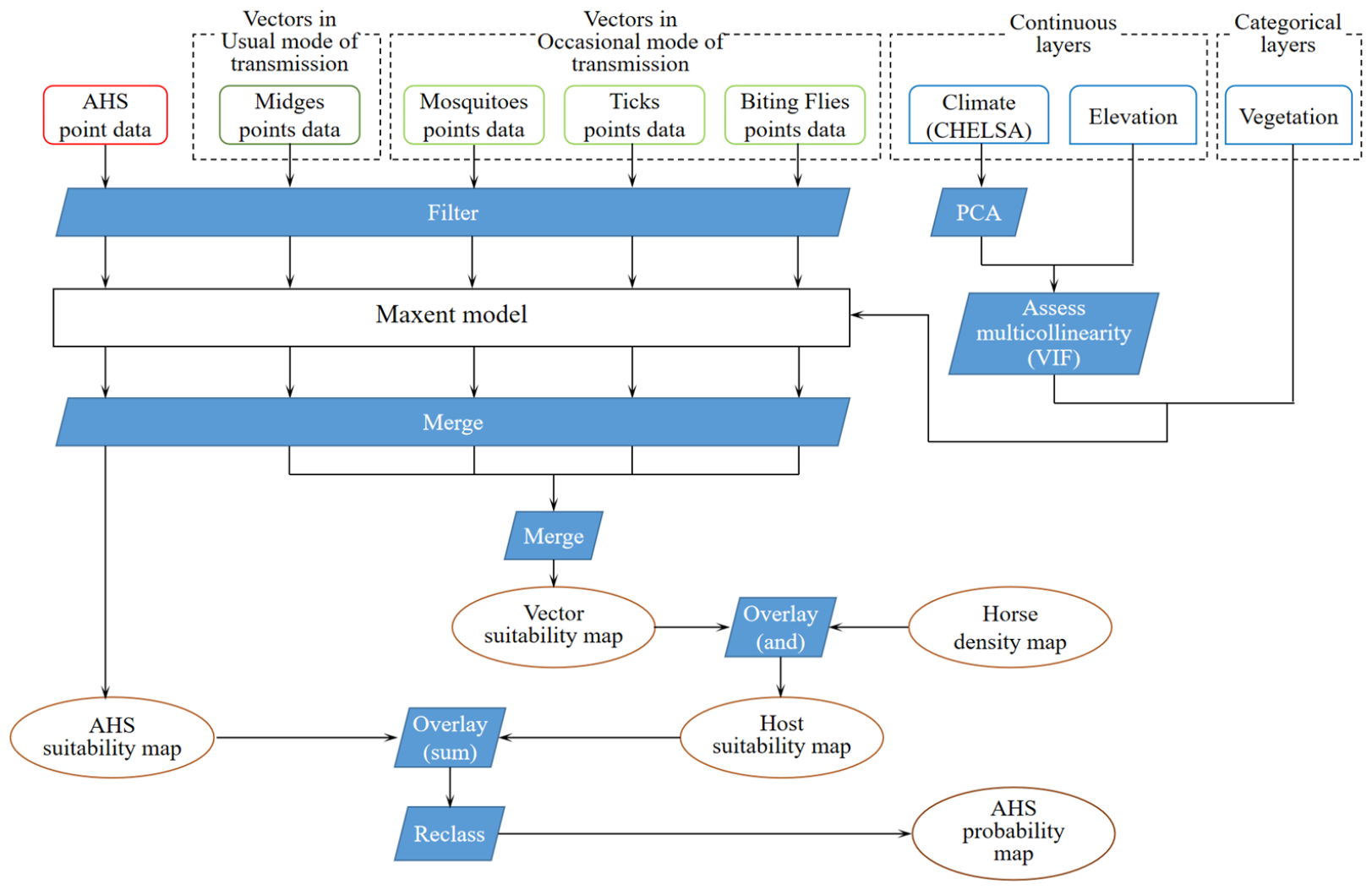
Flow Chart of the Modeling Process for AHS. (This figure presents a comprehensive flow chart outlining the step-by-step process of the AHS modeling. It includes stages such as data collection, processing, model development, and analysis, providing a clear visual representation of the methodology employed in the study.)

## 3. Results

### 3.1. Results of Data collection

From our comprehensive data collection efforts spanning 2005 to 2022, we identified 217 occurrences of ASH. Of these, 199 instances were recorded in Africa and 18 in Asia (details provided in Supplementary File 1). Our data compilation included extensive records of various vector species: 1,307 midges (including *C. imicola, C. bolitinos, C. variipennis*, and *C. brevitarsis*), 21,677 mosquitoes (*Aedes aegypti, Anopheles stephensi*, and *Culex pipiens*), 1,005 ticks (*Hyalomma dromedarii* and *Rhipicephalus sanguineus*), and 22,821 biting flies (*Stomoxys calcitrans* and *Tabanus* spp.), as detailed in Supplementary File 2.

### 3.2. Results of Climate-Based Zonal Segregation

Our analysis led to the creation of a climate classification map, categorizing regions into five groups: tropical, arid, temperate, continental, and polar/alpine (see Fig. 2). This classification resulted in a total of 20 distinct prediction units based on geographical regions and climate zones. Specifically, Africa comprises 3 units (tropical, arid, and temperate), Asia has 5 units (encompassing all five climate categories), Australasia is represented by 3 units (tropical, arid, and temperate), Europe by 4 units (excluding the arid category), and the Americas by 5 units (including all climate categories).

**Fig. 2.**
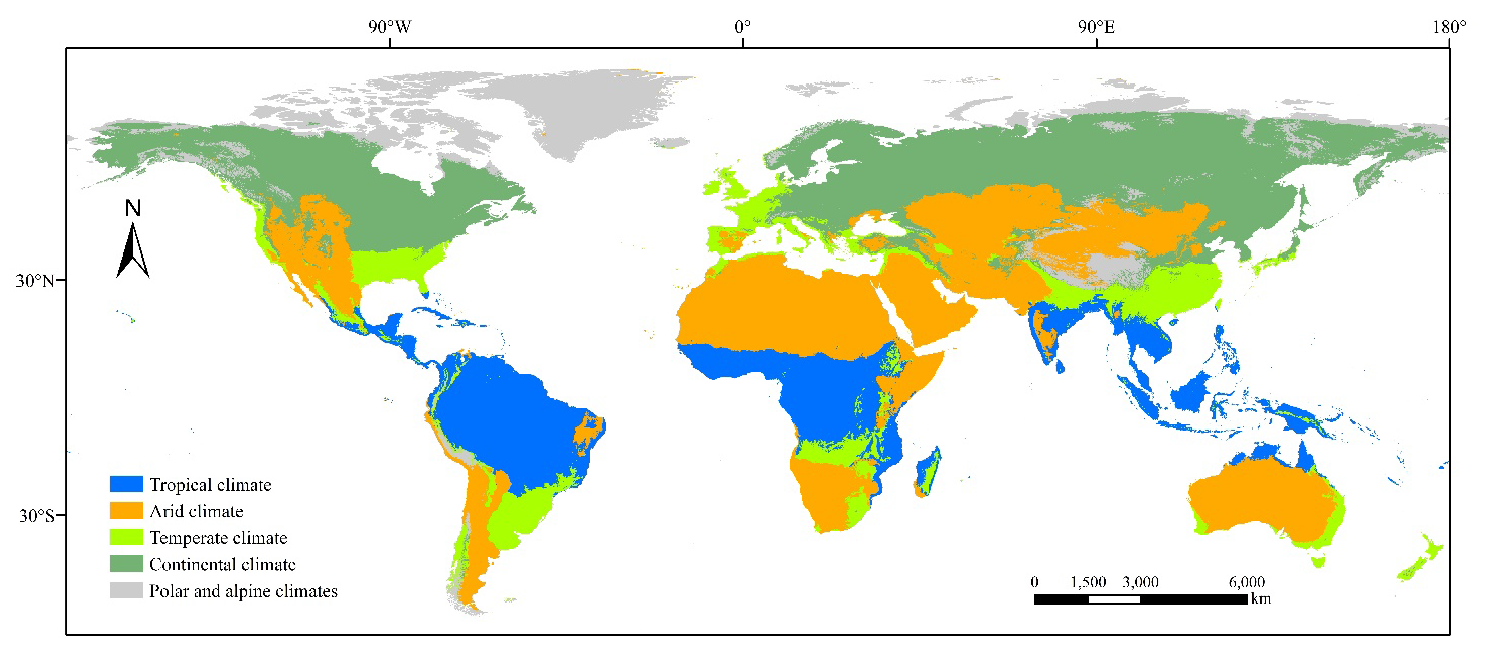
Climate Pattern-Based Classification of the Study Area. (This figure depicts the classification of the study area according to distinct climate patterns. It visually represents how the study regions are categorized into various climate zones, such as tropical, arid, temperate, continental, and polar/alpine, aiding in the understanding of the geographical and environmental context of the research.)

### 3.3. Modeling Outcomes for AHS Distribution

#### 3.3.1. Data Post-Filtering

Post data refinement, we identified 116 presence points of AHS. The modeling was conducted in various African regions (tropical, arid, and temperate) and tropical Asia. This allowed us to analyze AHS distribution in these diverse climatic zones.

#### 3.3.2. Key Predictors in Different Regions

The model identified region-specific key predictors for AHS distribution:

Tropical Africa: Land Cover, Elevation, and September mean air temperature (temp 9) emerged as primary predictors.

Tropical Asia: Key predictors included Land Cover, February mean air temperature (temp 2), February precipitation (prec 2), and July precipitation (prec 7).

African Arid Region: In this region, Land Cover, Elevation, October mean air temperature (temp 10), and September maximum air temperature (tmax 9) were significant.

African Temperate Region: Here, the model highlighted Land Cover, Elevation, precipitation of the driest month (bio 14), and February precipitation (prec 2) as crucial factors.

#### 3.3.3. Model Reliability and Variable Contribution

The VIF values for predictor variables ranged from 1.000 to 4.551, satisfying the low multicollinearity prerequisite (<10). The AUC values varied between 0.941 and 0.986, demonstrating a high degree of model reliability. Table 1 in the manuscript details the contribution rates, AUC values, and VIF values of the main variables incorporated into the model. Additionally, the response curves for each model are illustrated in Fig. 3. For ease of reference, an abbreviations table for the climate variables is provided in Supplementary File 3.

**Table 1.** Contribution Rates of Predictors and AUC Values in the AHS Model.

| Region | LC. | Elev. | Climate Variables |  |  |  |  |  |  | AUC | VIF |
| --- | --- | --- | --- | --- | --- | --- | --- | --- | --- | --- | --- |
|  |  |  | temp2 | temp9 | temp10 | tmax9 | bio14 | prec2 | prec7 |  |  |
| 1_Africa | 49.3 | 23.5 |  | 27.2 |  |  |  |  |  | 0.941 | 1.000-1.000 |
| 1_Asia | 29.4 |  | 32.8 |  |  |  |  | 24.3 | 13.5 | 0.951 | 1.074-1.526 |
| 2_Africa | 14.0 | 48.1 |  |  | 12.3 | 25.6 |  |  |  | 0.986 | 1.011-4.551 |
| 3_Africa | 11.3 | 32.2 |  |  |  |  | 40.1 | 16.4 |  | 0.982 | 1.052-3.509 |

1: Tropical Climate region, 2: Arid Climate region, 3: Temperate Climate region
LC.: Land cover, Elev.: Elevation

**Fig. 3.**
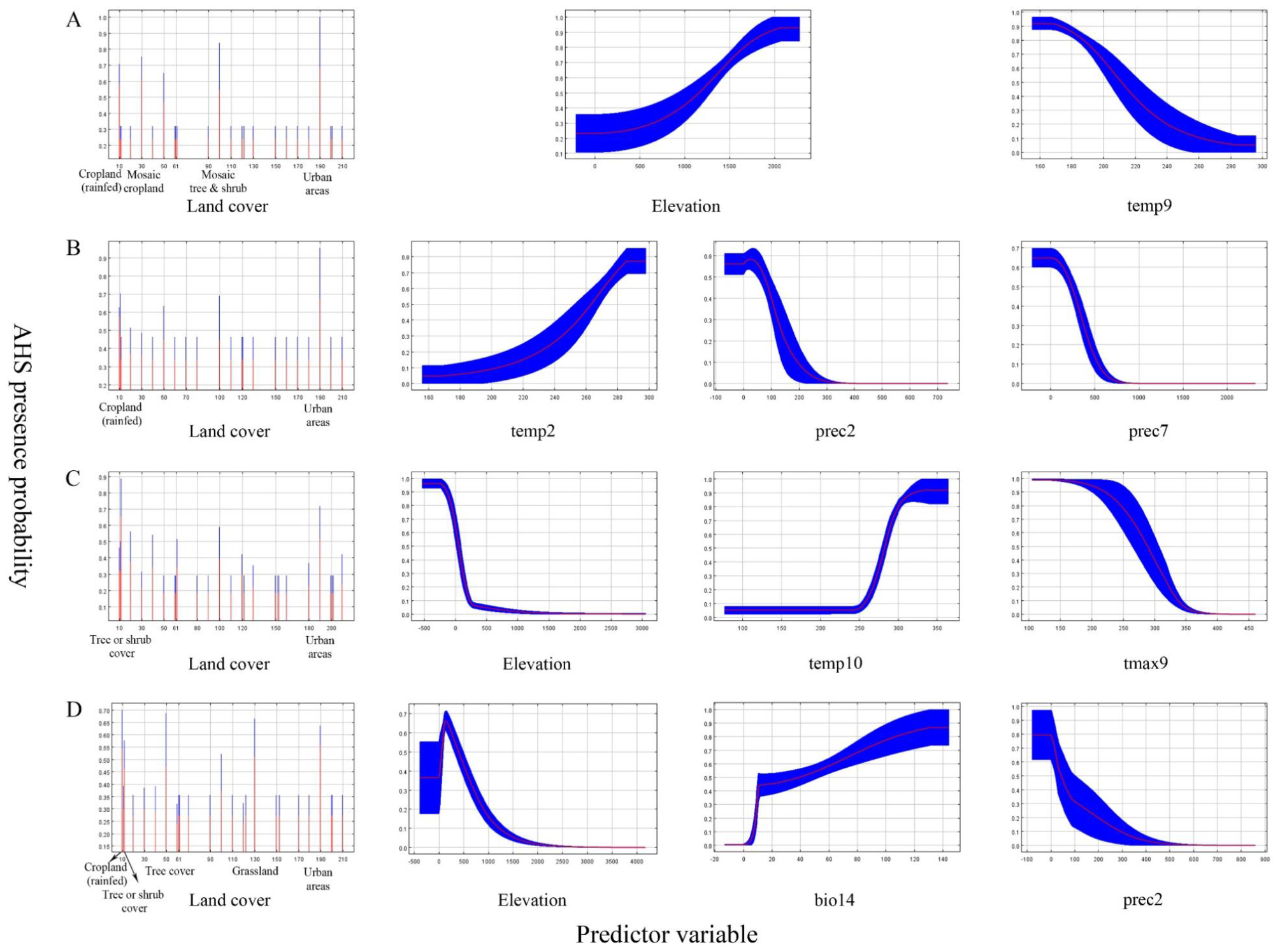
Response Curves of the AHS Model in Various Climate Regions. (This figure illustrates the response curves for the AHS model across different climate regions. The panels represent: (A) the Tropical Climate region of Africa, (B) the Tropical Climate region of Asia, (C) the Arid Climate region of Africa, and (D) the Temperate Climate region of Africa. Each curve displays the mean response (in red) and the associated mean standard deviation (in blue), providing insights into the model’s sensitivity to environmental variables in each region.)

### 3.4. Modeling Outcomes for AHSV Vector Distribution

#### 3.4.1. Data Post-Filtering

After meticulous filtering, our dataset encompassed presence points for various AHSV vectors: 1,229 midges, 9,751 mosquitoes, 796 ticks, and 10,574 biting flies. This comprehensive dataset facilitated the development of a total of 82 individual models tailored for niche modeling of AHSV vectors. These comprised 16 models for midges, 27 for mosquitoes, 15 for ticks, and 24 for biting flies.

#### 3.4.2. Model Robustness and Key Predictors

The VIF values for the predictors in these models ranged from 1.000 to 6.795, indicating low multicollinearity and hence, reliability of the models (<10 VIF threshold). The AUC values varied from 0.820 to 0.997, reflecting the robustness of the models. The primary environmental variables influencing vector distribution are detailed in Tables 2, 3, 4, and 5. The response curves for each vector model are available in Supplementary File 4.

**Table 2.**
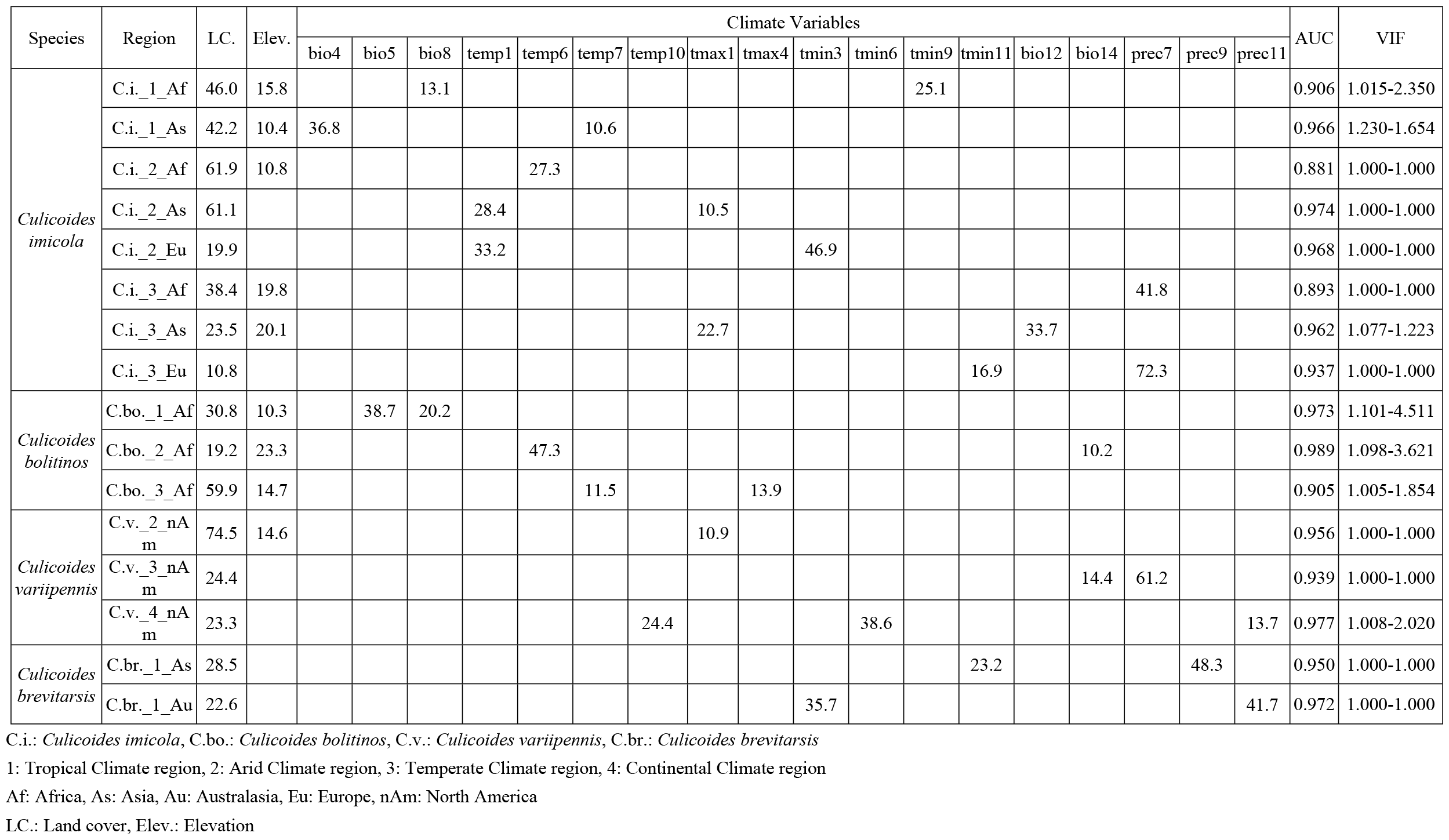
Contribution Rates of Predictors and AUC Values in the Distribution Models for *C. imicola, C. bolitinos, C. variipennis*, and *C. brevitarsis*.

**Table 3.**
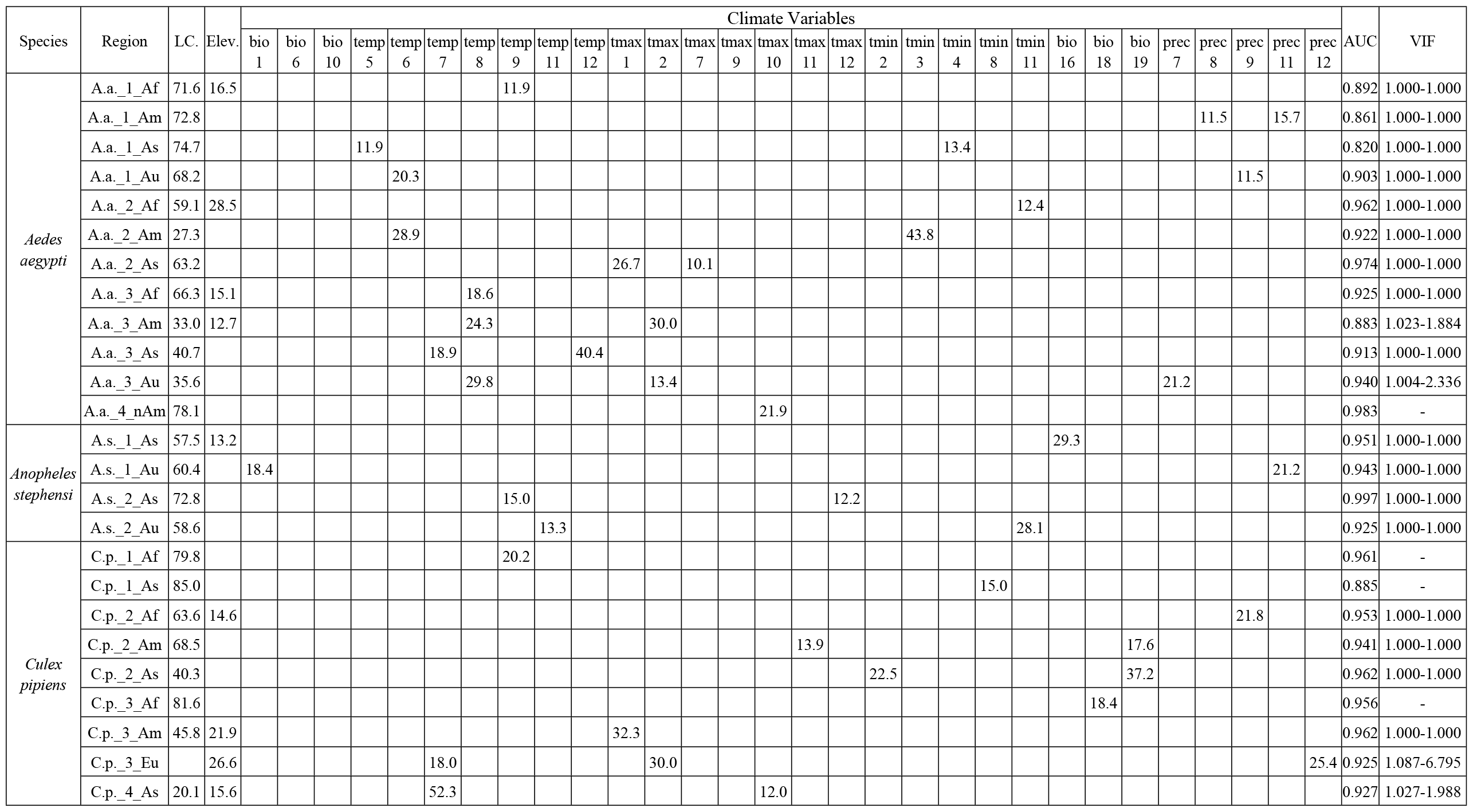
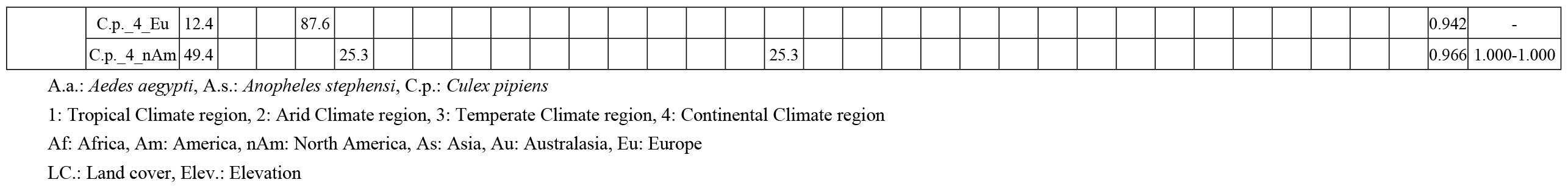
Contribution Rates of Predictors and AUC Values in the Distribution Models for *Aedes aegypti, Anopheles stephensi*, and *Culex pipiens*.

**Table 4.** Contribution Rates of Predictors and AUC Values in the Distribution Models for *Hyalomma dromedarii* and *Rhipicephalus sanguineus*.

| Species | Region | LC. | Elev. | Climate Variables |  |  |  |  |  |  |  |  |  |  |  |  |  |  |  | AUC | VIF |
| --- | --- | --- | --- | --- | --- | --- | --- | --- | --- | --- | --- | --- | --- | --- | --- | --- | --- | --- | --- | --- | --- |
|  |  |  |  | bio9 | bio11 | temp1 | temp6 | temp7 | temp9 | temp11 | tmax1 | tmax6 | tmax8 | tmin1 | tmin2 | prec6 | prec7 | prec8 | prec12 |  |  |
| *Hyalomma dromedarii* | H.d._2_As | 36.5 |  |  |  | 38.1 |  |  |  |  |  |  |  |  |  |  | 25.4 |  |  | 0.979 | 1.000-1.000 |
| *Rhipicephalus sanguineus* | R.s._1_Af | 87.2 | 12.8 |  |  |  |  |  |  |  |  |  |  |  |  |  |  |  |  | 0.864 | - |
|  | R.s._1_Am | 84.0 |  |  |  |  |  |  |  | 16.0 |  |  |  |  |  |  |  |  |  | 0.934 | - |
|  | R.s._1_As | 83.9 |  | 16.1 |  |  |  |  |  |  |  |  |  |  |  |  |  |  |  | 0.904 | - |
|  | R.s._2_Af | 74.1 |  |  |  |  | 15.1 |  |  |  |  | 10.8 |  |  |  |  |  |  |  | 0.975 | 1.000-1.000 |
|  | R.s._2_Am | 65.7 |  |  | 15.9 |  |  | 18.4 |  |  |  |  |  |  |  |  |  |  |  | 0.949 | 1.000-1.000 |
|  | R.s._2_As | 49.2 |  |  |  |  |  |  |  |  | 39.6 |  |  | 11.2 |  |  |  |  |  | 0.916 | 1.000-1.000 |
|  | R.s._2_Au | 51.4 |  |  |  |  | 28.7 |  |  |  |  |  |  |  | 19.9 |  |  |  |  | 0.922 | 1.000-1.000 |
|  | R.s._3_Af | 49.5 | 28.6 |  |  |  |  |  |  |  | 11.4 |  |  |  |  | 10.5 |  |  |  | 0.930 | 1.022-1.963 |
|  | R.s._3_Am | 81.9 |  |  |  |  |  |  | 18.1 |  |  |  |  |  |  |  |  |  |  | 0.922 | - |
|  | R.s._3_As | 55.5 |  |  |  |  |  |  | 20.7 |  |  |  |  |  |  |  | 23.8 |  |  | 0.933 | 1.000-1.000 |
|  | R.s._3_Au | 74.3 |  |  |  |  |  |  |  |  |  |  |  |  |  |  |  | 25.7 |  | 0.956 | - |
|  | R.s._3_Eu | 61.1 |  |  |  |  |  |  |  |  |  |  | 16.0 |  |  |  |  |  | 22.9 | 0.903 | 1.000-1.000 |
|  | R.s._4_Eu | 83.8 |  |  |  |  |  |  |  |  |  |  |  |  | 16.2 |  |  |  |  | 0.936 | - |
|  | R.s._4_nAm | 83.3 |  |  |  |  |  |  |  |  |  |  |  |  |  |  |  |  | 16.7 | 0.986 | - |

H.d.: *Hyalomma dromedarii*, R.s.: *Rhipicephalus sanguineus*
1: Tropical Climate region, 2: Arid Climate region, 3: Temperate Climate region, 4: Continental Climate region
Af: Africa, Am: America, nAm: North America, As: Asia, Au: Australasia, Eu: Europe
LC.: Land cover, Elev.: Elevation

**Table 5.**
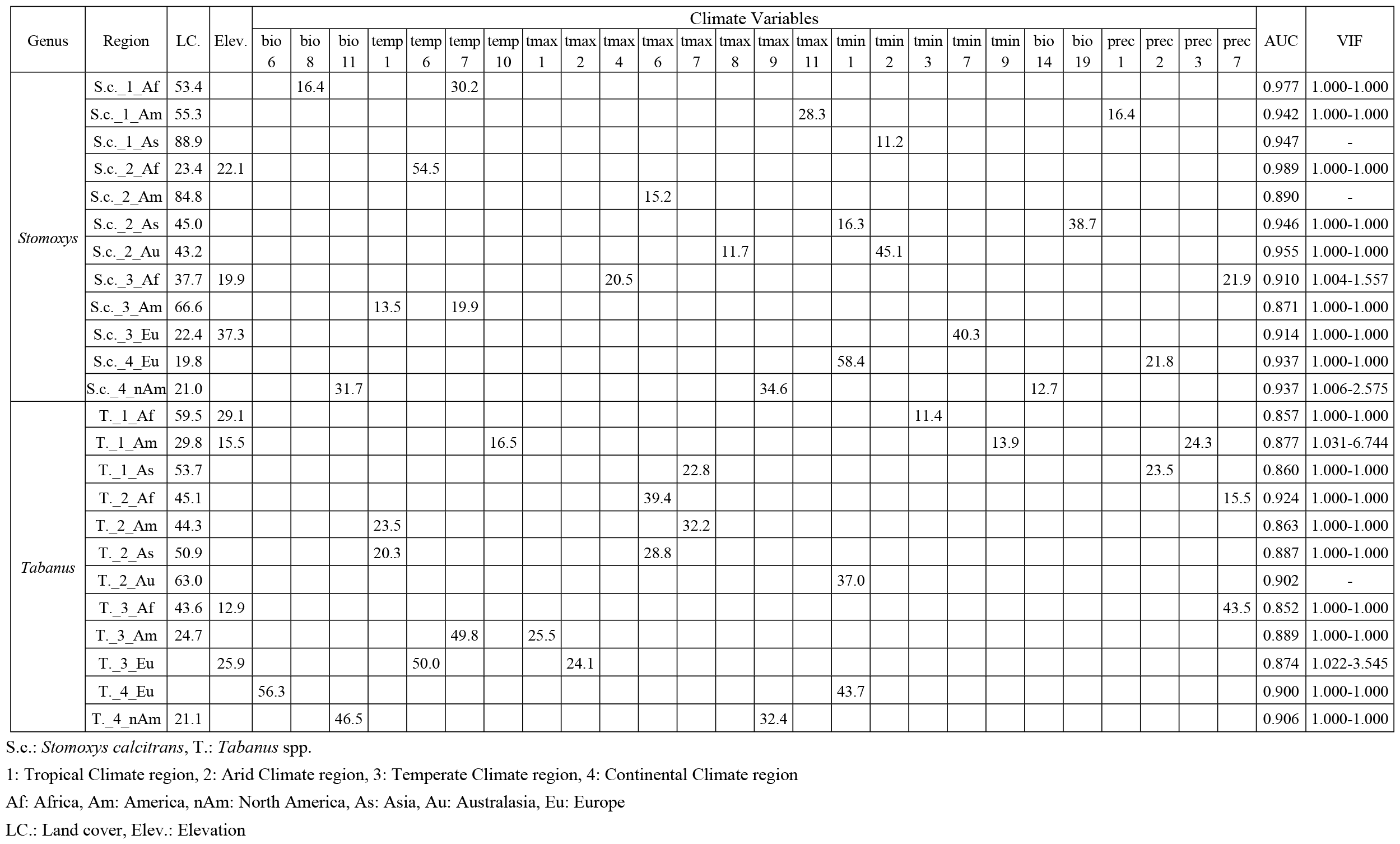
Contribution Rates of Predictors and AUC Values in the Distribution Models for *Stomoxys calcitrans* and *Tabanus* spp.

### 3.5. Identifying High-Risk Zones for AHS by Integrated Modeling Approach

Our integrated modeling approach pinpointed high-risk zones for AHS across the globe. Notably, a significant portion of Europe, excluding its northern areas, was identified as high-risk. In Asia, parts of the Indian subcontinent, South Asia, Southeast Asia, and East Asia, along with some regions of the Middle East, showed elevated risk levels. For Africa, the model highlighted the middle and southern regions as particularly vulnerable. In the Americas, extensive areas of the United States, Central America, and specific regions in South America, such as Brazil, were marked as high-risk zones. These findings are visually represented in Fig. 4.

**Fig. 4.**
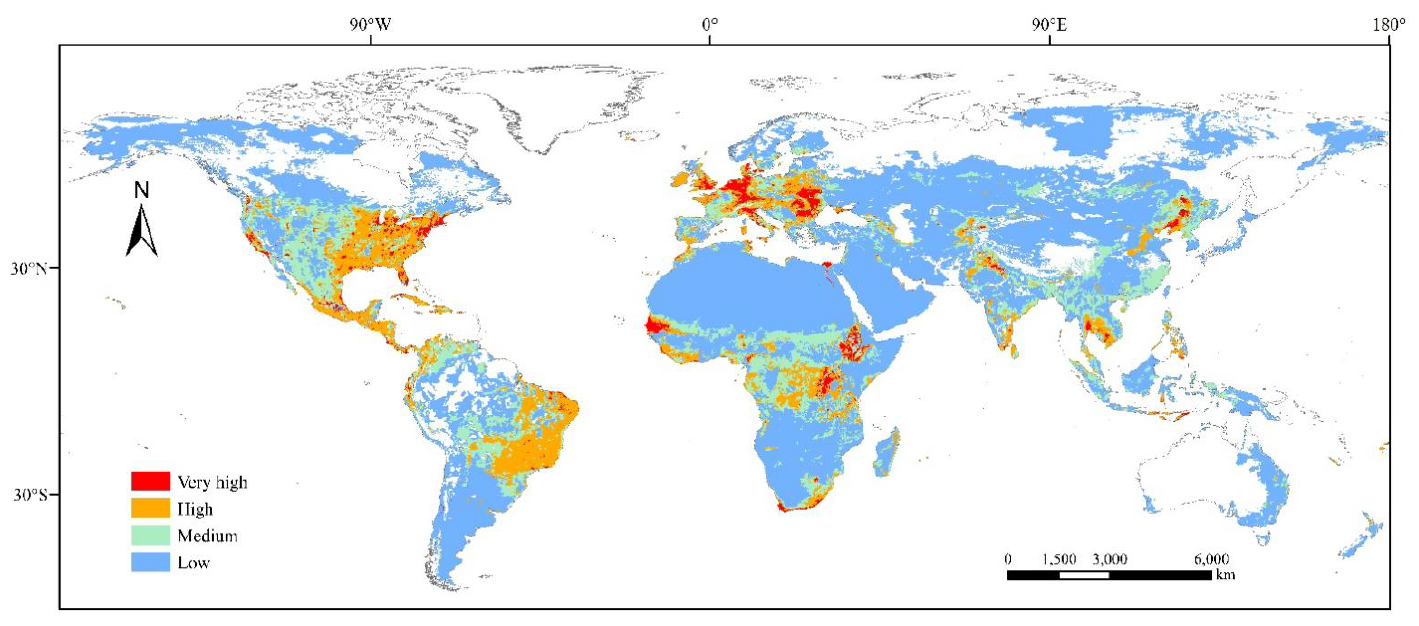
Global Distribution of High-Risk Zones for AHS.

## Discussion

Our research offers a novel perspective on the environmental factors influencing AHS in diverse climatic regions. We have identified key environmental variables that correlate significantly with AHS occurrences in both Africa’s varied climates and Asia’s tropical regions. This comprehensive approach not only substantiates the reliability of our identified factors but also aligns our findings with other region-specific prediction studies, thereby bolstering the credibility of our model outcomes. The research by Shan Gao et al. and Danica Liebenberg et al., which emphasizes factors like annual temperature ranges and precipitation, resonates with our results, especially regarding precipitation during crucial months and temperature variations (17, 42). These similarities validate our results and highlight the complex interplay between environmental factors, vertebrate hosts, and vector arthropods of AHSV, echoing previous research (44–48).

In recognizing the limitations of our study, it’s important to note the exclusion of certain equines, including wild species and donkeys, and the reliance on standard climate data. This limitation arises from the challenges in data availability and the difficulty of monitoring wild animal populations (49). The significant role of donkeys, known to be potential carriers or reservoirs of AHS due to their large population size and wide distribution, cannot be overlooked (50). Excluding donkey data might lead to an underestimation of risk areas or a failure to fully capture the disease’s transmission dynamics. Future predictions, therefore, require reassessment with more comprehensive and reliable data on donkeys and other wild equines when such data become available.

In our study, we relied on standard climate data rather than specific microclimate data, acknowledging the limitations this approach may impose. Microclimate data often provide more accurate predictions for the spread of vector-borne diseases (51), as they account for the specific environmental preferences of vector arthropods like Culicoides spp., a known AHSV vector (52–55). These vectors select resting areas that offer favorable conditions, significantly influencing disease transmission dynamics (56). The utilization of microclimate data is essential but also challenging, as it requires access to detailed environmental variables (57, 58). Our results, therefore, might overlook certain low-risk areas that actually harbor vectors, leading to discrepancies in risk assessment. Future research should aim to include more specific, reliable, and comprehensive data, emphasizing the need for advancements in data collection methodologies.

Our research plays a crucial role in shaping control and management strategies for AHS in high-risk areas. Our study emphasizes the importance of implementing a comprehensive approach that includes vector control, immunization, and public awareness. Additionally, we highlight the potential impact of climate change in intensifying AHSV vector habitats. Our findings underscore the need for future studies to incorporate more detailed environmental data to better understand and mitigate AHS risks.

The increasing prevalence of AHS is a significant environmental concern, largely attributed to climate change. Rising temperatures are expanding the habitats of AHSV vectors, notably *C. imicola*, leading to a northward spread and enhanced AHSV transmission (1, 12, 28, 59, 60). This change, coupled with the identification of new vector species, suggests an increased risk of AHS in broader areas (61, 62). Recent outbreaks of Bluetongue virus, transmitted by the same vectors, further highlight the growing concern about AHSV expansion (63, 64). Our findings emphasize the urgent need to consider these environmental changes in AHS management strategies.

In conclusion, our research contributes significantly to understanding AHS and its environmental drivers. Future studies should aim to integrate broader data sets, including microclimatic variables, and consider socio-economic factors for a more comprehensive approach. Collaboration with various stakeholders is vital for the practical application of our findings in disease management and control.

## Acknowledgments

I would like to extend my heartfelt appreciation to the participants of this study for their willingness to share their time, experiences, and insights. Their contributions have been invaluable in enriching the findings of this research.

## Notes

### Competing Interest Statement

The authors have declared no competing interest.

